# iCSD can produce spurious results in dense electrode arrays

**DOI:** 10.1101/2025.05.02.651822

**Authors:** Joseph Tharayil, Esra Neufeld, Michael Reimann

## Abstract

Estimation of the current source density (CSD) is a commonly-used method to interpret local field potential (LFP) signals by estimating the location of the neural sinks and sources of current that give rise to the LFP. We show analytically that, when the inter-electrode spacing is small relative to the width of the current distribution, commonly used methods to estimate the CSD produces spurious results, calculating true sources as sinks and vice versa. By simulating a biologically-detailed subvolume of the rat somatosensory cortex with over 200,000 biophysically-detailed neurons, we show that, for high-density recording electrodes, the estimated CSD diverges from expected results, necessitating awareness and careful interpretation of CSD results.

## 1 Introduction

The electrical potential signal recorded by electrodes in the extracellular space is generated primarily by neuronal transmembrane currents; the local field potential (LFP), the low-frequency component of these signals, is largely driven by synaptic currents [1]. The distribution of these currents in space is known as the current source density (CSD). Under quasi-static conditions, which generally hold for physiological environments, the CSD is related to the potential field in the tissue by Poisson’s equation [2].

Several methods have been developed to estimate the CSD from recordings made by electrode arrays [3, 4, 5, 6, 7, 8, 9]. In this paper, we focus on the “standard” CSD (sCSD) method [3] and the commonly-used inverse CSD (iCSD) method [4], which estimate the CSD from laminar electrode array recordings, assuming that the source distribution is homogeneous in planes perpendicular to the electrode array (or homogeneous within a specified distance and negligible beyond it, in the case of iCSD). However, in various brain regions, neural activity has been shown to vary on shorter (sub-millimeter) spatial scales [10, 11]. The iCSD has been found to be robust to inhomogeneity in the current distribution for a number of specified spatial CSD profiles [4]; however, it is not clear whether this robustness holds under realistic physiological conditions and for recently developed high density electrode arrays.

Simulations of biophysical models of neural circuits have been used to investigate the biological basis of the CSD signal [12]. In addition, because biophysical models permit the calculation of a ground-truth current source density, they have been used to investigate how accurately CSD estimation methods reconstruct the ground truth. In [13], Pettersen et al. use a model of 1000 Layer 5 pyramidal cells with passive dendrites to calculate ground-truth and estimated current source densities in response to a simulated whisker flick stimulus. In [14], Hagen et al. use a model of 4000 Layer 4 spiny stellate cells and 1000 Layer 4 Large Basket Cells, both with passive dendrites, and with both populations receiving thalamic, but not recurrent, synaptic input. In both [13] and [14], iCSD methods have been found to better estimate ground-truth current distributions than the standard CSD method. Others have used biophysical modeling to compare ground-truth data with CSD estimates produced using the 2D-iCSD [5] and kCSD [15] approaches. However, previous studies have neural models with limited morphological and electrical diversity, and which include only feed-forward, and not recurrent, connectivity. As CSD is derived from the LFP, which is thought to be primarily synaptically-driven, one important source of spatial inhomogeneities in true CSD is the co-activation of excitatory and inhibitory synapses on the same dendrites. A model without recurrent connectivity, including inhibitory connectivity, cannot capture this.

Our analytical investigations have suggested that such inhomogeneities might be particularly problematic for high-density electrode arrays, such as the recently developed Neuropixels probes [16, 17]. In this paper, we investigate this issue by simulating LFP recordings from the Blue Brain Project (BBP) model of the rat somatosensory cortex [18, 19]. We calculate the CSD from these recordings using the sCSD and iCSD methods, and compare them to the “objective CSD” metric derived from the model ground truth. We specifically focus on how the density of electrode contacts affects the agreement between objective CSD and the estimated CSDs.

## 2 Methods

### 2.1 Analytic investigation

Under quasi-static conditions, the current source density *C* is related to the extracellular potential *ϕ* by Poisson’s equation:

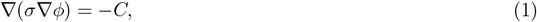

where *σ* is the tissue conductivity and the electric field *E* = −∇*ϕ*.

Under the assumption that the tissue is homogeneous and isotropic, and that *ϕ* is constant in the x-y plane (perpendicular to the electrode array), Eq 1 simplifies [4] to

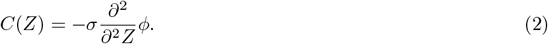

Evaluating a discretized version of Equation 2, namely

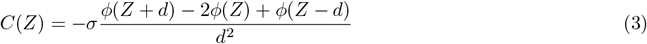

where *d* is the inter-electrode spacing, is the “standard” CSD method.

For a point source current with magnitude *I*, the potential *ϕ* at a point at a distance *x* is given by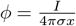. We denote the prefactor 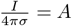 for simplicity.

For a laminar electrode array with inter-electrode spacing *d*, the potential at a particular electrode caused by a point source current at a radial distance *r* and a vertical distance *z* is given by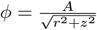. By Equation 2, the contribution *C*_*p*_ to the estimated CSD (after discretization according to the recording resolution, and under the assumption of homogeneity in the planar direction) of this point source is

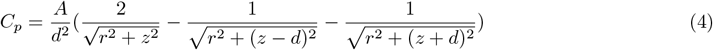

Substituting 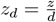 and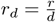, Eq 4 evaluates to

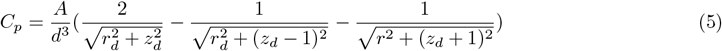

As a function of *z*_*d*_, the density of points such that *C*_*p*_ = *p* for arbitrary *p* is given by

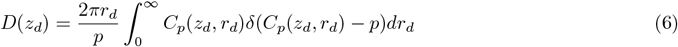

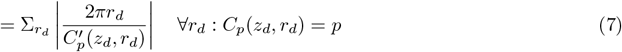

where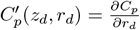. We solve this equation numerically over a range of *z*_*d*_ for selected values of *p*.

If, instead of a point source, the current distribution is a 2-dimensional Gaussian with width *w* at depth distance *z* and centered on the electrode array, the contribution *C*_*g*_ to the CSD is given by

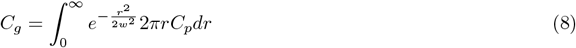

Substituting 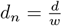 and 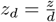, Eq. 8 evaluates to

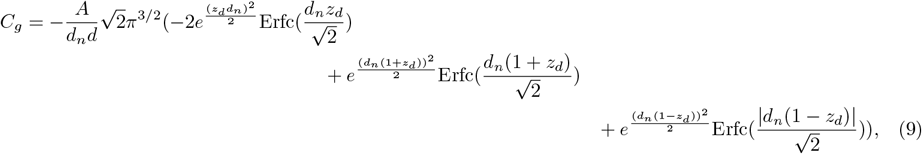

where Erfc is the complementary error function; 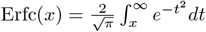

In the limit of homogeneous activity in the x-y-plane, we obtain:

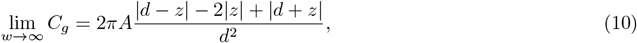

which is equal to 0 when | *z*| ≥ *d* and to 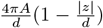 otherwise. We are also interested in the limit case of an infinitely fine electrode array. Defining *z*_*n*_ = *z/w*,

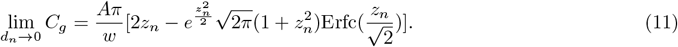

### 2.2 iCSD

Inverse CSD (iCSD) is a commonly used technique to estimate current distributions, using less restrictive assumptions than the standard CSD technique [4]. In this work, we use the step-iCSD variant of the iCSD technique, which assumes that the current distribution is homogeneous within a cylinder with height *h* (typically equal to the array spacing) and radius *ρ* (estimated based on the expected spatial extent of the relevant current sources).

### 2.3 Biophysical models

We simulate a 7-column subvolume of the BBP rat somatosensory cortex model [18]. The circuit consists of *∼*200,000 multicompartmental neuron models, where each compartment represents a small discretized segment of a neurite. For each of the *k* compartments in the circuit, a transmembrane current `i_membrane` is computed at each time step. Ornstein-Uhlenbeck conductance noise is injected in order to simulate missing synaptic inputs. To account for the fact that applying spike sorting algorithms to *in vivo* recording systematically underestimates firing rates [20, 21, 22], noise input is optimized to produce *in silico* firing rates equal to 30% of recorded *in vivo* firing rates (*P*_*FR*_ = 0.3), with the ratio of of the standard deviation of the noise to the mean of the noise fixed at 0.4 (*R*_*OU*_ = 0.4)[19]. A virtual whisker flick stimulus is applied with 10% of virtual projections from the ventral posteromedial nucleus activated, as in [19], for 10 trials. Simulations are conducted with a simulated extracellular calcium concentration of 1.05 mM.

### 2.4 In Silico LFP and CSD

We simulate LFP recordings from laminar electrode arrays in the biophysical model. LFP signals are calculated using the line-source approximation option of the BlueRecording software suite [23]. For each electrode *k* and each neural compartment *j* in the model, BlueRecording calculates a coefficient *w*_*k,j*_, such that *LFP* (*k, t*) = Σ_*j*_*w*_*k,j*_`i_membrane`_*j*_(*t*), and writes the values of *w* to a “weights file”, which is read by the Neurodamus simulator to calculate the LFP at simulation run time. For more details, see [23]. Extracellular conductivity is set to 0.375 S/m; as conductivity is assumed to be homogeneous, this acts as a simple scaling factor. The CSD is calculated using the step-iCSD method [4], as implemented in [12], for various values of *ρ*.

### 2.5 Non-negative CSD

We define the “non-negative CSD” (nnCSD) as the sum of the compartmental contributions to the CSD according to the standard CSD method, but excluding compartments that contribute to the signal with the opposite sign of their associated current (negative sensitivity). To calculate nnCSD, we take the negative second finite difference with respect to *Z* of the weights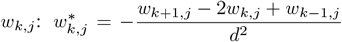. tThe standard CSD estimate (sCSD) can be computed as 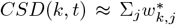 `i_membrane`_*j*_(*t*) (because the CSD is approximately the second finite difference of the LFP, and the calculation of the LFP is linear). The nnCSD is obtained as 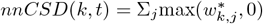`i_membrane`_*j*_(*t*).

### 2.6 Objective CSD

We define an “objective disk” current source density *o*_*D*_*CSD* (Fig. 1a), at a given time step and electrode. A current source contributes to the *o*_*D*_*CSD* at electrode *k* if it is within a disk perpendicular to the electrode array, centered on electrode *k* and with a thickness *t* equal, by default, to the inter-electrode spacing and a radius *ρ* equal, by default, to 500 *µ*m:

**Figure 1.**
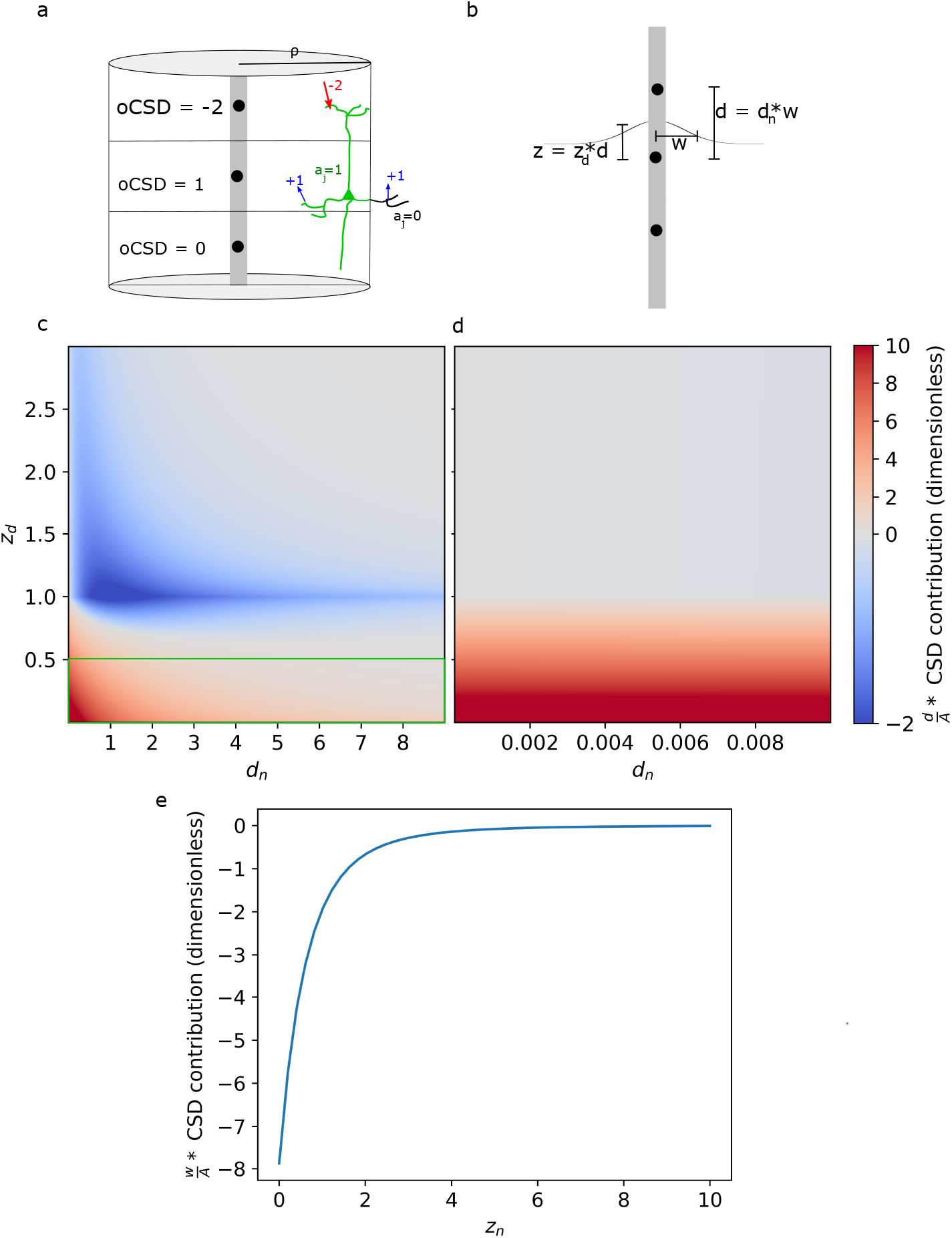
a: Illustration of the *o*_*D*_*CSD* calculation. The coefficients *a*_*j*_ for each neural segment are represented by the color of the segment. Arrows represent transmembrane currents, with magnitudes specified by adjacent numbers. *o*_*D*_*CSD* is the sum of all currents within a disk of radius *ρ* centered on each electrode. b: Gaussian current source distribution centered on the electrode array. c: Normalized contribution to the CSD from a Gaussian current distribution centered on the electrode, as a function of *d*_*n*_, the inter-electrode spacing normalized by the width of the Gaussian and *z*_*d*_, the difference in depth of the source and the electrode-of interest, normalized by *d*. We cancel out the prefactor 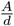 in Eq. 9, as we are primarily interested in the sign of the contribution. The green box indicates the region which, for the *o*_*D*_*CSD*, would have a positive CSD contribution, while the rest of the space doesn’t contribute to *o*_*D*_*CSD*. d: Same as Panel a, but zoomed on very small values of *d*_*n*_. e: Normalized CSD contribution as a function of *z*_*n*_, in the limit as *d*_*n*_ goes to zero, and for finite *w*. We cancel out the prefactor 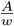 in Equation 11.

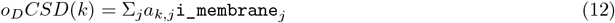

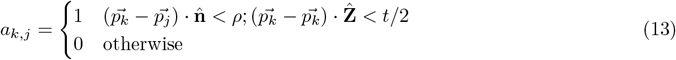

where 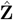 refers to the unit vector in the direction of the electrode axis, perpendicular to the plane of the disk, and 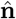 refers to the unit vector normal to the electrode axis, in the plane of the disk. We note that because *o*_*D*_*CSD* is not normalized by the volume of the disk, it is a measure of total current rather than current density. Therefore, when comparing *o*_*D*_*CSD* measures obtained using different parameters, and when comparing *o*_*D*_*CSD* with other metrics, all measures are first normalized by their respective root-mean-squared amplitudes.

*o*_*D*_*CSD* is equivalent to the measure that the step-iCSD method [4] aims to estimate for the same electrode layout and with radius *ρ*. That is, if the assumption of the iCSD method of uniformity within the disk is true, then both measures will yield the same result up to a constant scaling factor (accounting for the volume of the disk). The *o*_*D*_*CSD* computation is implemented in BlueRecording. The *o*_*D*_*CSD* is similar to the ground-source CSD metric defined in [14]; however, the metric in [14] accounts for segments that are partially inside and partially outside the disk by weighting their contributions to *o*_*D*_*CSD* by the fraction of their lengths inside the disk. In contrast, our method fully includes the contribution of segments with centers within the disk, and ignores the contribution of segments with centers outside the disk.

### 2.7 Implementation and code availability

- Code for running the simulations and generating the figures in this paper is available at https://github.com/joseph-tharayil/csd_paper. Postprocessed LFP traces from these simulations are available on Zenodo under the following doi: https://doi.org/10.5281/zenodo.14998743.
- The source code for BlueRecording is available at https://github.com/BlueBrain/BlueRecording.
- The seven column subvolume of the BBP circuit model is available under the following DOI: `10.5281/zenodo.7930276`.

Simulations are performed using the Neurodamus [24] simulation control application, with the CORENEURON [25] computation engine. LFP and *o*_*D*_*CSD* signals were recorded at 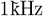 sampling rate; we analyzed a 50 ms window beginning at the onset of the whisker flick stimulus. The neural simulations described in this paper were run on the BB5 supercomputer using 32 nodes; each node has two 2.30 GHz, 18 core Xeon SkyLake 6140 CPUs, and 382 GB DRAM.

## 3 Results

### 3.1 Dense electrode arrays may result in confusion of sinks and sources

Using Equation 9, we calculate the contribution to the estimated, discretized standard CSD, of a 2-D Gaussian current source centered on the array, as a function of *d*_*n*_, the inter-electrode spacing normalized by the width of the Gaussian, and *z*_*d*_, the vertical distance from the electrode to the center of the Gaussian, normalized by the inter-electrode distance (Fig. 1b). We visualize the quantity 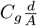, canceling out the prefactor 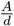 in Equation 9, as we are primarily interested in the sign of the contribution and canceling 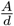 permits us to express the contribution as a function of only two, dimensionless parameters. Note that the standard CSD and iCSD methods assume uniformity over at least part of the horizontal plane [4], while the Gaussian current source is not uniform in that plane. Thus, we investigate the impact of violating the assumption of in-plane uniformity, to assess under which circumstances non-uniform current sources in biological tissue will adversely affect the iCSD estimates.

For all values of *d*_*n*_, the contribution to the CSD will be negative for *z*_*d*_ *∼* 1 (that is, for a Gaussian centered on the neighboring electrode; Fig. 1c). However, as the electrode array becomes more dense, the range of values of *z*_*d*_ producing negative CSD contributions increases. The analytically-calculated CSD contribution differs from the *o*_*D*_*CSD*, in which the contribution from a Gaussian centered on the electrode array would be positive for *z*_*d*_< 0.5, and 0 otherwise, for all values of *d*_*n*_.

While decreasing *d*_*n*_ is associated with a greater range of *z*_*d*_ which produce negative contributions, this does not hold for *d*_*n*_ *<<* 1. In this regime, realistic widths of the Gaussian are much larger than the electrode spacing. Hence, this case approximates uniformity in the horizontal plane, i.e. where the assumption underlying iCSD methods is true. Given that in this regime the assumptions underlying iCSD are true, we do not expect to see any confusion of sources and sinks. As expected, when *d*_*n*_ is very small (*d*_*n*_ *<* 0.01) we found no negative CSD contributions (Fig. 1d).

However, it is unlikely that *in vivo* recording setups are in this ideal regime. For an array with an electrode spacing of 50 *µ*m, a Gaussian current distribution would need to be *∼* 3.9mm wide to avoid negative weights below −0.1. (In comparison, the radius of a cortical column is *∼* 200*µm*.) It is thus unlikely that the limit case of an effectively homogeneous current distribution actually occurs *in vivo*. Therefore, for a given current distribution at an unknown depth, as the recording electrode array density increases, and consequently *d*_*n*_ decreases, the probability that the current distribution produces a spurious CSD contribution (i.e., a source where there should be a sink, or vice versa) increases. *In vivo*, the CSD is produced by a large number of currents distributed throughout the tissue. Thus, our analysis indicates that, for dense electrode arrays, at least some of these currents are likely to produce spurious CSD contributions.

For an infinitely fine electrode array, the CSD contribution from a Gaussian current source with finite width is given by Eq. 11. We visualize the quantity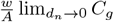, since canceling out the prefactor 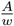 permits us to express the contribution as a function of only one, dimensionless parameter. We find that a such a current source with will produce negative contributions to the CSD at effectively all locations (Fig 1e).

We next attempt to determine if violation of the assumption of homogeneity is a practical issue in our model. For an array with inter-electrode spacing of 20*µ*m, we calculate *C*_*p*_ as a function of radius and *z*_*d*_ (Fig. 2a, left). We observe a positive lobe centered on *z*_*d*_ = 0 and flanked by two negative lobes. We calculate the densities *D*_*p*_ for two isocontours at *C*_*p*_ = −.003 and *C*_*p*_ = .004. Both have minimal density at *z*_*d*_ = 0. For the positive isocontour, the density is concentrated in a narrow region of *z*_*d*_, while it increases more gradually for the negative isocontour (Fig. 2a, right).

**Figure 2.**
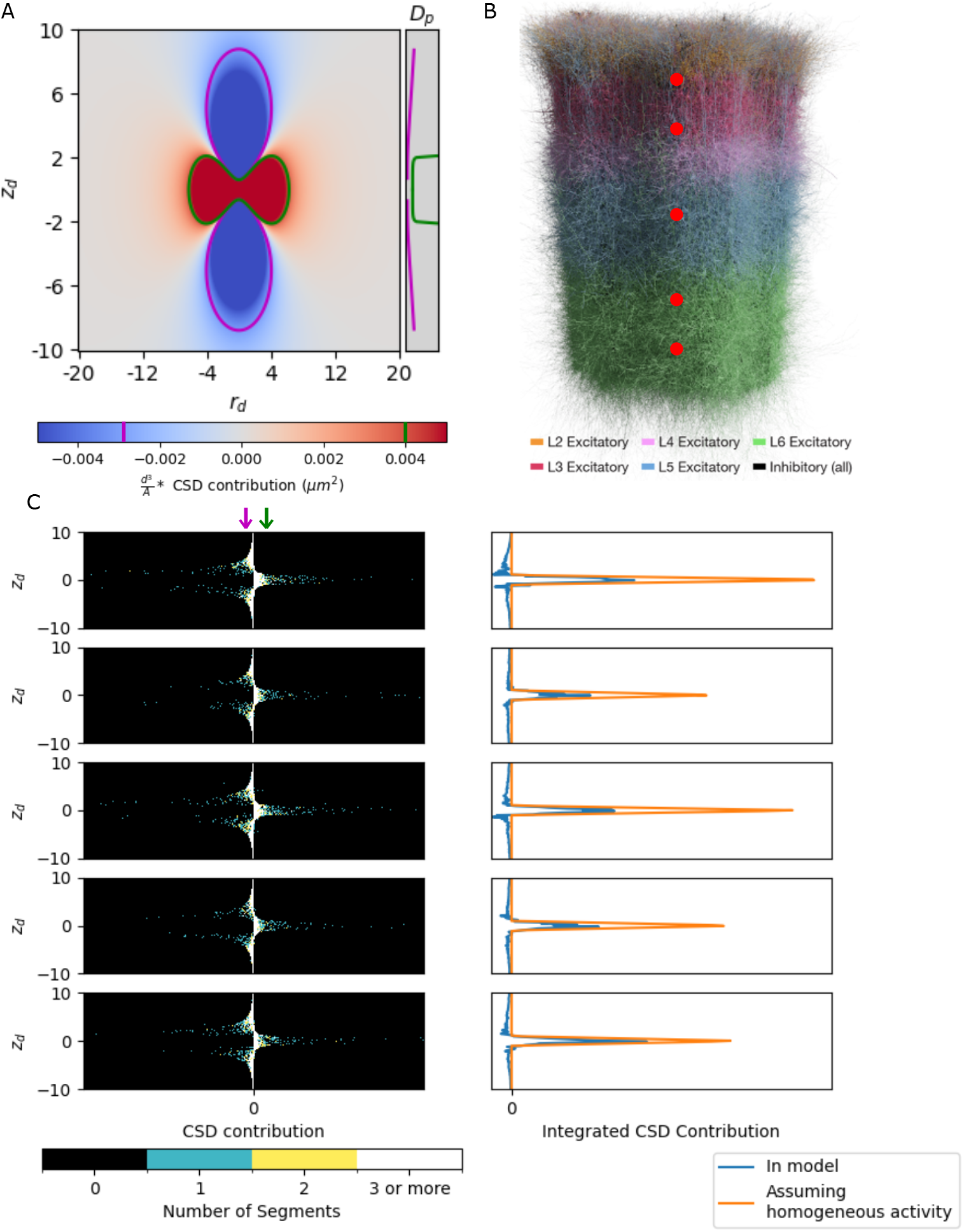
A: Left: *C*_*p*_ as a function of radius and *z*_*d*_, for a value of *d* = 20*µm*. Green and purple lines represent selected positive and negative isocontours, respectively. Values of the isocontours are indicated by vertical lines on the colorbar. Right: Density *D*_*p*_ for the isocontours in the left panel, as a function of *z*_*d*_. B: The cortical columm. Red circles represent the positions of the electrodes used in Panel C. C: Left column: Histogram of CSD contributions from a unit current at each neural segment, as a function of the segment’s vertical distance from the electrode. Bin size in the *z*_*d*_ direction is 0.177, and 10^−8^ in the *C*_*p*_ direction. The CSD contribution levels corresponding to the isocontours in panel A are indicated by green and purple arrows. We note that the density of positive contributions is concentrated around *z*_*d*_ = 0, as observed in Panel A, and that this is flanked by the density of the negative contributions. Right column: Integrated CSD contribution from all segments, as a function of vertical distance from the electrode (blue trace) and the analytic integrated CSD contribution assuming a homogeneous, continuously distributed source, scaled to match the total absolute CSD contribution from the model. Each row represents a different electrode, ordered from the top of the column to the bottom.

We then calculate *w*_*k,j*_, the contribution of a unit current from the *j*-th neural compartment to the LFP recorded at the *k*-th electrode, for an array of 101 electrodes centered on the cortical column, and with an inter-electrode spacing of 20 *µ*m. We then take the negative second finite difference of the contributions with respect to the axis of the electrode array, in order to obtain the contributions 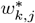 of a unit current at each neural compartment to the CSD recorded at the *k*-th electrode.

For five different electrode contacts (at 200 *µ*m, 500 *µ*m, 1000 *µ*, 1500 *µ*, and 1800 *µ* from the bottom of the column, Fig. 2b) and for neural segments from a randomly sampled 5% of neurons in the central column, we find subsets of compartments with strongly positive weights 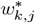 around the contact depth (i.e., | *z*_*d*_|< 1), and with large negative values in the depth-range 1 <|*z*_*d*_|< 5) (Fig. 2c). The large number of negative 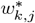 clearly demonstrates that the CSD sign flipping issue is not a purely theoretical problem. The problem persists even in the ideal limit of a homogeneous current distribution in the x-y plane: We would expect no contribution to the CSD at *z*_*d*_ ≥ 1 (Eq. 10, Fig. 1d). However, we find that there are big depth ranges where the sum of CSD contributions is negative (Fig. 2c). This must logically be a result of the discrete nature of the neuron morphologies, which do not continuously fill the space – an effect that particularly matters in electrode proximity, where signal contributions are high and the distance-dependent potential varies rapidly due to nonlinearity. Effectively, the negative deflections in Fig 2c can be explained by sampling in the negative lobes observed in Fig 2a, while the positive deflection results from sampling from the positive lobe. Thus, even if all of the neural segments in a plane had identical transmembrane currents, the measured CSD would not perfectly match the analytic expectation and can negatively contribute to recordings at depth-distances *d*.

### 3.2 iCSD diverges from true current source density

Estimation methods such as iCSD make weaker assumptions about homogeneity in the horizontal plane. To further explore our hypothesis and test whether confusion of sinks and sources is a relevant issue for iCSD in realistic models, we performed *in vivo*-like simulations of a whisker flick stimulation protocol (see Methods), modeling laminar electrode arrays with inter-electrode spacings of 160, 80, 40, and 20 *µ*m, and calculate the iCSD and *o*_*D*_*CSD*. At array densities of 160 *µ*m and 80 *µ*m, the iCSD is almost identical to the *o*_*D*_*CSD* (Fig 3 a.i vs a.v; a.ii vs a.vi). At an array density of 40 *µ*m, the iCSD (Fig 3 a.iii) begins to diverge from the *o*_*D*_*CSD* (Fig 3 a.vii), displaying high-frequency spatial oscillations in the iCSD that are not observed in the *o*_*D*_*CSD*. This effect becomes even more pronounced at an array density of 20 *µ*m (Fig 3 a.iv vs a.viii).

**Figure 3.**
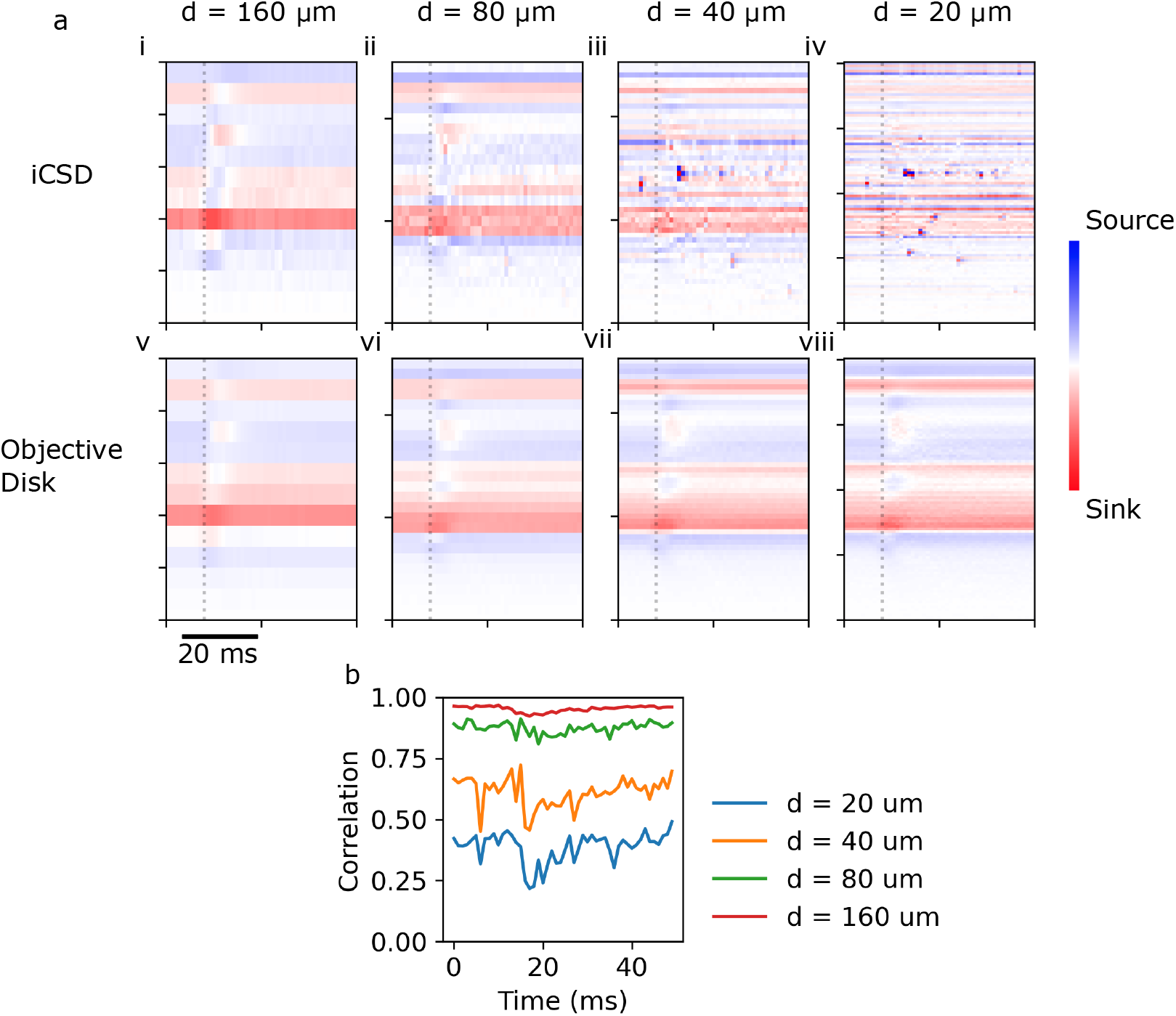
iCSD diverges from *o*_*D*_*CSD* (calculated with radius *ρ* = 500*µm*) at high array densities. a: iCSD (top row) and *o*_*D*_*CSD* (bottom row) for array densities of 160 *µ*m, 80 *µ*m, 40 *µ*m, and 20 *µ*m (first through fourth columns, respectively). The whisker flick is initiated at *t* = 0, and arrives in the cortex at time *t ∼* 10ms (grey dashed line). b: Correlation over the depth profile, as a function of time, between iCSD and *o*_*D*_*CSD*, for various electrode arrays.

We calculated the Pearson correlation over the depth profile, at each time point, between the iCSD and *o*_*D*_*CSD*, for each electrode array. While the two metrics are highly correlated for the 160 *µ*m and 80 *µ*m arrays, the correlation decreases for the 40 *µ*m array.(Fig. 3 b). For the 20 *µ*m array, the correlation is much lower throughout the time course of the signal compared to those of the other arrays (Fig. 3 b).

### 3.3 Error is due to inhomogeneity in true current distribution

Given the results of our analytical investigation (Fig.1a), it is natural to interpret the high-frequency spatial noise in the iCSD (Fig 3a.iii,iv) as the result of current sources appearing as sinks and vice versa. However, there is an alternative explanation for the results observed so far. It may be that iCSD, when applied to recordings from very dense electrode arrays, is able to resolve very high (spatial) frequency fluctuations that are actually present in the current density, but are not detected by coarser electrode arrays. Because we used *o*_*D*_*CSD* with a constant radius parameter of *ρ* = 500*µ*m, the *o*_*D*_*CSD* averages currents over a large radius, and is likely to smooth over such fluctuations. If this latter interpretation is correct, we would expect that the discrepancy between iCSD and *o*_*D*_*CSD* would reduce when reducing the current averaging radius *ρ*.

Therefore, we compare the iCSD, nnCSD (a metric based on standard CSD, but which excludes the contributions of neural segments in locations where sources and sinks are confused; see Section 2.5 for details), and *o*_*D*_*CSD* calculated for various disk radii, for an electrode spacing of 20 *µ*m (Fig. 4a). We observe that for *ρ* = 100*µ*m and *ρ* = 50*µ*m, the correlation between iCSD and *o*_*D*_*CSD* is substantially better than for larger values of *ρ* (Fig. 4b). One interpretation of this result is that, with decreasing *ρ*, high-spatial-frequency oscillations in the current distribution close to the electrode cease to be averaged out. However, this does not explain why the correlation between iCSD and *o*_*D*_*CSD* drops for *ρ* = 20*µ*m. An alternative interpretation is that part of the discrepancy between iCSD and *o*_*D*_*CSD* is caused by misspecification of the radius over which the current distribution is assumed to be constant; and that at *ρ* = 100*µ*m, this error is minimized. Under this interpretation, the oscillations themselves, which are still visible in the iCSD at *ρ* = 100*µ*m, are attributable to confusion of current sinks and sources – the oscillations remain because even over a radius of 100*µm*, the underlying current distribution is not entirely homogeneous. In support of this latter hypothesis, the correlation between nnCSD and *o*_*D*_*CSD* is higher than the correlation between iCSD and *o*_*D*_*CSD* for all radii (Fig. 4d). Increased misspecification for low *ρ* would also explain the emergence of a pronounced late current source in iCSD in the middle of the column at *ρ* = 20*µ*m (Fig. 4a): This might be explained as another inversion of sources and sinks. Furthermore, we note that the pattern commonly of interest in such an experiment, i.e., the laminar distributions of sinks and sources triggered by the whisker flick, is most evident in the *o*_*D*_*CSD* for *ρ* = 200*µm*, where the correlation with iCSD is lowest.

**Figure 4.**
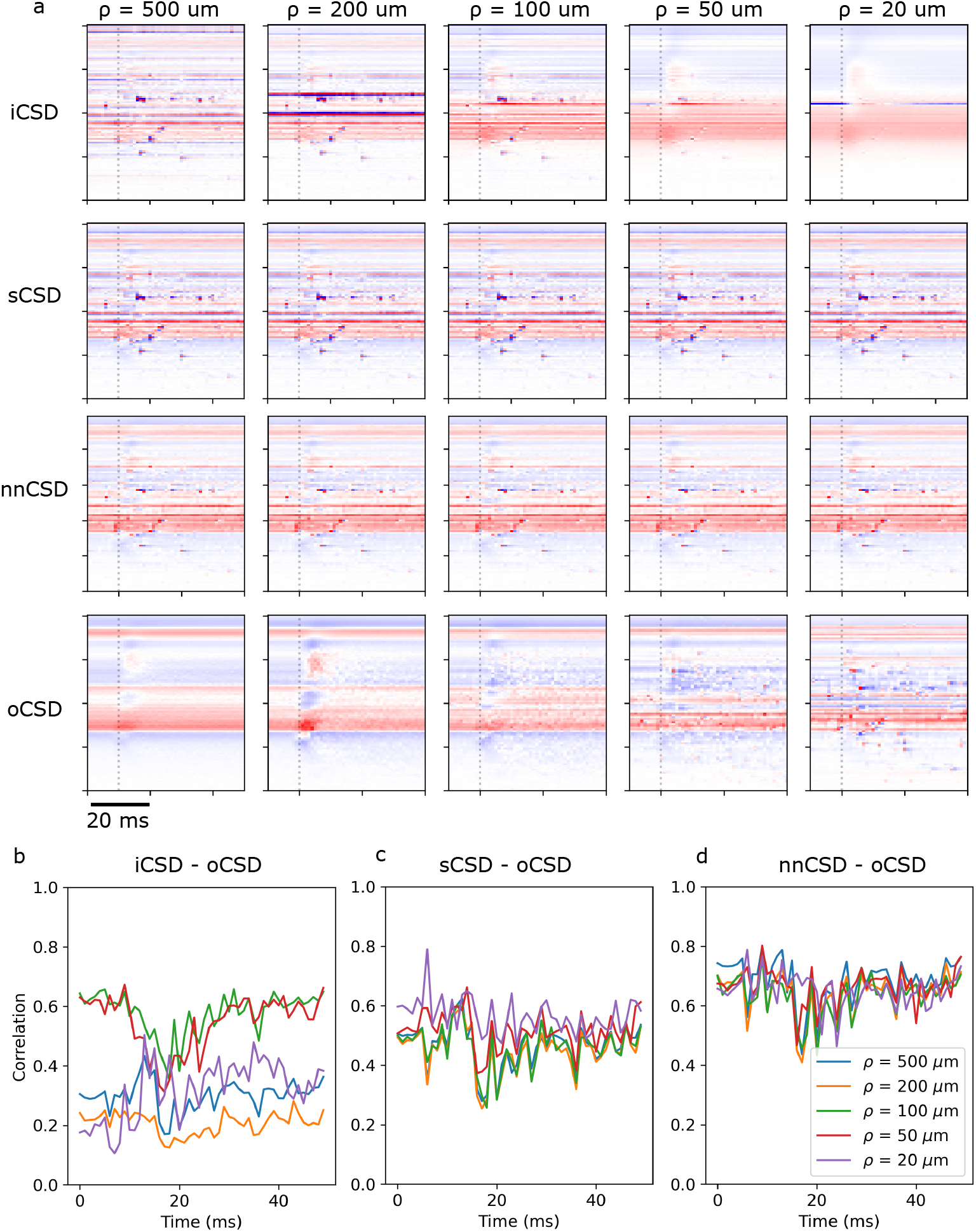
a. iCSD, sCSD, nnCSD, and *o*_*D*_*CSD* recorded for a 20 *µ*m electrode spacing. The radius *ρ* for *o*_*D*_*CSD* and iCSD is varied. The whisker flick is initiated at time *t* = 0, and arrives in the cortex at time *t ∼* 10ms (grey dashed line). b: Correlation between iCSD and *o*_*D*_*CSD* with various cylinder radii. c: Correlation between sCSD and *o*_*D*_*CSD* with various cylinder radii. d: Correlation between nnCSD and *o*_*D*_*CSD* for various cylinder radii.

However, it is not necessarily the case that the improvement in the iCSD-*o*_*D*_*CSD* correlation due to decreasing *ρ* can be attributed entirely to reduction in the inhomogeneity of the current distribution; it may be that more accurate calculation of true oscillations simultaneously plays a role. Given that high-frequency spatial oscillations can be seen in the iCSD for *ρ* = 500*µ*m, we also expect these oscillations to occur in the standard CSD. We therefore repeat the investigation performed above, using standard CSD instead of iCSD (Fig. 4a). If the observed high-frequency oscillations are real, but averaged out in *o*_*D*_*CSD*, we expect to see an increased correlation between sCSD and *o*_*D*_*CSD* for decreased *ρ*. Because sCSD, unlike iCSD, is not parameterized by *ρ*, improvements in the sCSD-*o*_*D*_*CSD* correlation cannot be explained by reduced error in the assumption regarding the radius over which the current is homogeneous. Therefore if the sCSD-*o*_*D*_*CSD* correlation does increase with decreasing *ρ*, this would indicate that the high-frequency oscillations are a true feature of the current distribution.

We observe that the correlation between sCSD and *o*_*D*_*CSD* does not change substantially as *ρ* is varied (Fig. 4c), and that nnCSD consistently correlates better with *o*_*D*_*CSD* than does iCSD (Fig. 4d). Together, these results suggest that for the 20 *µ*m electrode spacing, discrepancies between estimated CSD and *o*_*D*_*CSD* are primarily due to confusion of sinks and sources due to an inhomogeneous underlying current distribution, and not to improved resolution of true sinks and sources.

## 4 Discussion

We have shown analytically that, for small inter-electrode spacings, sCSD may produce spurious results when the assumption of in-plane homogeneity in the ground-truth current distribution does not hold (Fig. 1c). We further show that the potential for confusion of sinks and sources increases as array density increases, and that in our model, compartment weights can be strongly negative and CSD would differ from the analytical expectation (becoming negative at a depth distance > *d*) even if in-plane currents were identical, because neural trajectories sample the space with a discrete resolution (Fig. 2).

Moreover, we have demonstrated that in our circuit model, iCSD diverges from *o*_*D*_*CSD* for dense electrode arrays (Fig. 3). Because *o*_*D*_*CSD* is the ground-truth average of all current sources and sinks within a cylinder centered on the electrode, the difference between iCSD and *o*_*D*_*CSD* suggests that, for dense electrode arrays and under the conditions simulated in this paper, iCSD does not accurately reflect the ground-truth current distribution.

In principle, it may be the case that iCSD and *o*_*D*_*CSD* diverge because the iCSD from high-density electrodes captures high-spatial-frequency oscillations that are averaged out in *o*_*D*_*CSD*, particularly over large radii *ρ*. However, the correlation between nnCSD and *o*_*D*_*CSD* is consistently higher than the correlation between iCSD and *o*_*D*_*CSD* (Fig. 4b vs d), suggesting that confusion of sinks and sources likely accounts for the discrepancy between iCSD and *o*_*D*_*CSD*. In addition, we find no relationship between the value of *ρ* and the correlation between standard CSD and *o*_*D*_*CSD* (Fig. 4c), which indicates that *o*_*D*_*CSD* does not average out true high-spatial-frequency oscillations.

Previous studies using biophysical models have shown that iCSD [13, 14] does accurately reproduces ground-truth current distributions. However, [13], as well as several *in silico* experiments in [14], use an inter-electrode spacing of 100 *µ*m. As we have shown both analytically (Fig. 1) and numerically (Fig. 3), large inter-electrode spacings permit a more accurate reconstruction of the underlying CSD than do dense arrays. Still, [14] has shown that iCSD can accurately reconstruct ground-truth CSD even with a 20 *µ*m electrode spacing.

This may be because the neural circuit model in [14] produces a more uniform current distribution than the more detailed and realistic model studied here. Because the model in [14] is driven entirely by thalamic input, with no recurrent connectivity or extrinsic drive, the spatial scale of the current distribution is determined by the radius of the thalamic inputs. Conversely, in a model with recurrent connectivity, excitatory and inhibitory synapses are co-located on the same dendrites in close proximity. Unlike in a purely feedforward network, the co-activation of these excitatory and inhibitory recurrent synapses will result in an inhomogeneous CSD over small spatial scales. Thus, the recurrent connectivity in our model, coupled with a greater impact from active currents due to a more realistic level of intrinsic and evoked spiking activity, is expected to result in a current source distribution that is not truly homogeneous at any scale. In addition, the model in [14] includes only Layer 4 spiny stellate and basket cells, resulting in a much smaller spatial extent in the *z*-direction than our model. It may be that, simply due to the larger scale of our model, a greater number neural segments are present in regions (i.e., ranges of the parameter *z*_*d*_) where sinks and sources are confused (Fig. 1).

### 4.1 New interpretations for *in vivo* results

In [26], Zhang et al. observed that iCSD recorded in the macaque visual cortex using a laminar electrode with 20 *µ*m electrode spacing had a significant amount of noise along the depth axis, but that this noise disappeared after downsampling the electrode array or applying a Gaussian filter to the CSD. The authors surmise that the observed noise is due to nearly-balanced sinks and sources in thin bands of cortical tissue. However, these results may also be due to potential confounding of sinks and sources by iCSD, which we expect: If the spatial correlation in the CSD is driven by the combination of synaptic and return currents in individual neurons, rather than correlations between neurons, the larger size of primate neurons compared to rodent neurons would lead to a wider CSD, and therefore to a smaller value of *d*_*n*_ for a given electrode size (Fig. 1 a).

### 4.2 Outlook

We have shown that interpretation of iCSD obtained from the latest generation of laminar electrode arrays needs to be approached with caution. In particular, researchers should be aware of the possibility that iCSD from dense arrays confounds sinks and sources. Even if the high frequency noise in the iCSD (Fig.3a.iii,iv) does represent true fluctuations in the current distribution, their interpretation is not straightforward. Typically, CSD is used to investigate the large-scale architecture of entire pathways. However, the high (spatial) frequency component is likely sensitive to variability of synapse placements of individual axons. It thus is possible that iCSD in coarse arrays reflects trends in the activation of synapses over the entire dendritic tree, but that at higher resolution, iCSD resolves the synaptic currents, flanked by the associated return currents. In this case, iCSD calculated from high-resolution arrays would reflect the local electrical structure of invididual neurons, rather than the innervation pattern of the entire region.

Our results suggest that the assumptions made in CSD calculations ought to be revisited, particularly when using the latest generation of laminar electrodes. Rather than simply assuming uniformity in the current distribution in space, more realistic assumptions could be derived from simulation results, and the CSD calculation tuned accordingly.

In iCSD, accurate specification of the radius parameter *ρ*, over which the current distribution is assumed to be homogeneous, improves the agreement between the estimated and ground-truth current source distributions [14]. Previous work has shown that iCSD replicates the shape of the true current distribution even when the radius is misspecified by 50% [4]; even a misspecification by an order of magnitude results in a decrease in the correlation between ground-truth and estimated CSD of only *∼*10% [14], even for dense (20 *µ*m spacing) electrode arrays. We have shown that in our model, misspecification of radius can have a more dramatic effect; reduction of *ρ* from 200 to 100 *µ*m results in a factor *∼* 3 improvement in the correlation between iCSD and *o*_*D*_*CSD* (Fig. 4b). That misspecification of *ρ* has a greater effect in our model than in [14] is likely because of greater overall homogeneity in the ground-truth current distribution in the circuit model used in that paper, as discussed above.

*In silico* simulations may be useful in deriving more accurate assumptions about the homogeneity of the current sources. By calculating the *o*_*D*_*CSD* for a 2D grid of electrodes spanning the horizontal extent of the circuit model, it may be possible to estimate the effective width of the current distribution, which can in turn be used to approximate the value of *ρ* for the iCSD correction. Alternatively, *ρ* can be estimated from comparisons of *o*_*D*_*CSD* and iCSD; in our model, comparisons of *o*_*D*_*CSD* and iCSD indicated an ideal value of *ρ* between 50 and 100 *µ*m (Fig. 4b).

The use of alternatives to iCSD may permit a more accurate estimation of the current distribution when using dense electrode arrays. Methods such as 2D-iCSD [5] and kCSD [15], have previously been found to accurately estimate ground-truth current densities. While [5] and [15] used relatively coarse electrode arrays, kCSD makes fewer assumptions than iCSD regarding the underlying current distribution [6]. It is therefore possible that it will be less affected by inhomogeneity in the ground-truth CSD.

## Acknowledgments

This work was supported by funding to the Blue Brain Project, a research center of the École polytechnique fédérale de Lausanne (EPFL), from the Swiss government’s ETH Board of the Swiss Federal Institutes of Technology. We thank Prof. Gaute Einevoll for fruitful discussions regarding this paper.

